# Effect of germicidal short wave-length ultraviolet light on the polyphenols, vitamins, and microbial inactivation in highly opaque apple juice

**DOI:** 10.1101/2022.07.29.502038

**Authors:** Anita Scales Akwu, Ankit Patras, Brahmiah Pendyala, Anjali Kurup, Fur-Chi Chen, Matthew J. Vergne

## Abstract

The aim of this investigation was to study the efficacy of UV-C light emitting diode system (LED) operating at 263 nm for the inactivation of *Listeria monocytogenes* and *Escherichia coli* O157:H7. Specified concentrations of bacteria were inoculated in apple juice and irradiated at the designated UV doses of 0 to 15 mJ·cm^-2^. In addition, UV irradiation doses ranging from 0 to 160 mJ·cm^-2^ were also delivered to apple juice and polyphenols and vitamins were profiled. LC-MS/MS analysis was conducted to assess the stability of polyphenols or vitamins in UV-C exposed apple juice. The polyphenol and vitamin results demonstrated that UV-C irradiation in apple juices at relevant commercial UV doses induced significant reductions in the concentrations of selected polyphenols and vitamins, p<0.05. Ascorbic acid was reduced to 32%, at 160 mJ/cm^2^ whereas 17% reduction was observed at 40 mJ/cm^2^. Riboflavin was observed to be relatively stable. Epicatechin and chlorogenic was significantly reduced at high exposure doses. In contrast minor changes were observed at 40 mJ/cm^2^. Results show that UV-C irradiation effectively inactivated pathogenic microbes in apple juice. The log reduction kinetics of microorganisms followed log-linear and with higher R2 (>0.95) and low RMSE values. The D_10_ values of 4.16 and 3.84 mJ·cm^-2^ were obtained from the inactivation of *Escherichia coli*, and *Listeria monocytogenes* in apple juice. The results from this study imply that adequate log reduction of pathogens is achievable in apple juice and suggest significant potential for UV-C treatment of other liquid foods.

## Introduction

Apples and their based products are excellent sources of some important bioactive compounds, including key polyphenols responsible for their antioxidant properties (Kahle et al., 2005; Khanizadeh et al., 2008; Van der Sluis et al., 2002; Marcotte et al., 2022) including water and lipid soluble vitamins. Bioactive compounds present in fruits are significantly important to the human body due to its high antioxidant activity. In apples, polyphenols have demonstrated lung tissue damage prevention derived from individuals that smoke (Bao et al., 2013). Due to the presence of various flavonoids and other vitamins, apples have been extensively investigated as an important source of bioactive compounds (Boyer and Liu, 2004).

Phytochemicals are significantly important to the human body because of their antioxidant content, which works in preventing damage to our organ health. In apples, polyphenols have demonstrated lung tissue damage prevention derived from individuals that smoke (Bao et al., 2013). There are five major groups of polyphenols present in apples, to include, catechins, epicatechin, and procyanidins, quercetin, glycosides, phenolic acids (chlorgenic, gallic and coumaric acids), dihydrochalcones (phloretin glycosides), and anthyocanins (cyanidin) (Boyer and Liu, 2004; Kschonsek et al., 2018). Though polyphenols are present in apples, the amounts can diminish during the juicing process because of oxidative conditions, to include pulping, pressing, and clarification (Oszmianski et al., 2007; Markowski et al., 2009). Vitamin C is present in large quantities, alongside of fibers, and pectin in fruits prior to their processing. It is very important in processing that the products are safe for consumption, free of microbes and have their nutrient content retained.

In order to protect against possible vegetative cells and spores, most fruit juices are pasteurization or sterilized. FDA’s 1998 final rule (63 FR 37030) for processing fruit and/or vegetable juice requires processors to label products with a warning of potential illness upon consumption of the product unless the product was processed to inactivate the most likely of pathogenic bacteria in the product by 5-log. This rule made processors employ conventional heat treatments for pasteurization to ensure that a 5-log10 reduction has been achieved to avoid the use of a warning label. FDA recommends that retail processors who treat their juice follow these same practices for their 5-log reduction processes. In addition, to the juice labeling regulation, retail processors must follow all applicable state laws and regulations. Heat process negatively affects the nutritional quality of fruit juices including degraded sensory properties. Various authors have reported the effect of heat on vitamins and other polyphenols (Patras et al, 2020; Patras et al., 2011). Therefore, there is a need of alternative pasteurization and sterilization techniques.

The 2000 report published by United States Food and Drug Administration, Center for Food Safety and Applied Nutrition entitled “Kinetics of Microbial Inactivation for Alternative Food Processing Technologies: Executive Summary” clearly stated the need for additional research pertaining to the exploration and validation of alternative food processing technologies as a means of producing safe and nutritional dense foods. (CFSAN, 2000). Innovative and cutting-edge technologies will need to be validated through scientific research to provide “proof of principle” to ensure the effectiveness of the newer technologies. Nowadays, UV light processing is one of the most promising alternatives to thermal food preservation treatments (Islam et al., 2016b; Müller et al., 2014; Kaya et al., 2015; Keyser et al., 2008; Franz et al., 2009; Caminiti et al., 2012). This method of pasteurization has been successfully used for juices under the Code of Federal Register (CFR 179.39).

UV-C irradiation covers part of the electromagnetic spectrum from 100 to 400 nm and is categorized as UV-A (320 to 400 nm), UV-B (280 to 320 nm), and UV-C (200 to 280 nm). UV-C is most germicidal against most types of microorganisms since photons are absorbed by DNA. The photochemical reactions occurring in the DNA inhibits the microbial replication (Patras et al., 2020). The use of UV-C irradiation has been suggested for disinfection of the fluids using low pressure lamps and light emitting diodes (LED). Primarily there is huge interest on LED to be used in UV research. High optical output LED’s devices are available and have been tested by various authors (Kurup et al., 2022). There have been a vast number of studies using ultraviolet light emitting diodes (LEDs); however, two major common weaknesses are the lack of optical data in the studies (Caminiti et al., 2012; Unluturk et al., 2010) and the verification of UV dosage. These and other factors have led to considerable variability in reported inactivation kinetics. Therefore, there is an urgent need for a rigorous evaluation of the UV sensitivity of bacteria.

This research study attempted to address the shortcomings of earlier studies, delivering verified UV doses, and evaluating the effects of UV treatment on apple juice using LED devices. Specifically, this study evaluated the effect of UV-C irradiation on the vitamins and polyphenols of apple juice. In addition, this study also evaluated the ability of UV-C irradiation to inactivate *Listeria monocytogenes* and *Escherichia coli* O157:H7 apple juice.

## Materials and methods

### Chemicals

Phloridzin dihydrate, formic acid, and acetonitrile were bought from Sigma Aldrich, USA, whereas Chlorogenic acid was bought from Extrasynthese, France. (+)-Catechin and (-)- epicatechin were bought from Adooq Bioscience, CA, USA. Standard reference material (SRM) 3257 catechin calibration standard solutions set was a gift from the National Institute of Standards and Technology, MD, USA.

Furthermore, analytical grade riboflavin, thiamine hydrochloride, pyridoxal hydrochloride, pyridoxine, pyridoxamine dihydrochloride, cyanocobalamin, choline chloride, biotin, niacin, niacinamide, lumichrome, ammonium formate and formic acid were purchased from Sigma Aldrich, Missouri, USA. HPLC and LCMS grade water, methanol and acetonitrile were sourced from Fisher Scientific, USA.

### Sample Preparation

Apples (cv Gala) were obtained from a local grocery store. All apples were thoroughly washed and juiced using a Brentwood 800W juicer. The juice was strained and then filtered using Whatman (28-30 µm, 2-3 µm). Optical data was obtained and recorded and 45mL per culture tube were stored at -20°C wrapped with aluminum foil until further processing, to avoid exposure to light. Two consecutive filtration steps were carried out to remove solid particulates. Prior to irradiation, apple juice samples were randomly assigned to a treatment process and thawed at room temperature. Following the juicing process, baseline optical data was obtained, and apple were divided into 20 aliquots of 10 ml each stored at -20 °C until used. Physicochemical characteristics of the apple juice is shown in Table 1. A balanced design with three replicates randomized in experimental order was performed for each UV dose.

**Table 1.** Physiochemical characteristics of apple juice.

| Parameters | Values |
| --- | --- |
| pH | $3.3 \pm 0.164$ |
| Total soluble solids (%) | $14.3 \pm 0.018$ |
| Titrateable acidity | $5 \pm 0.406$ |

27 Data presented as mean ± stdev

### Bacterial strains and cultural conditions

Two non-pathogenic and non-outbreak strains of different bacteria were used in this study *Escherichia coli* O157:H7 (35150) and *Listeria monocytogenes* (19115). The bacterial strains were procured from American Type Culture Collection (ATCC). The bacterial cultures were stored in 25% glycerol in cryovials at -80 °C. Fresh bacterial suspensions were prepared for inoculation into apple juice for every treatment. Two loops of individual strains of *E. coli* and *L. monocytogenes* individually were transferred to 15 mL Tryptic soy broth (Oxoid Ltd., Basingstoke, UK) and incubated at 37 °C for 18 h. *L. monocytogenes* was also subjected to two successive transfers in tubes containing 15 mL of Buffered Listeria enrichment broth (Oxoid Ltd., Basingstoke, UK) and incubation was done for 24 h at 37 °C. These cultures were used as the adapted inoculum. After incubation, *E. coli* culture was transferred into 15 mL of TSB and incubated for 18 h at 37 °C to reach the stationary growth phase. Similarly, *L. monocytogenes* culture was transferred to 15 mL Listeria enrichment broth (Oxoid Ltd., Basingstoke, UK) and incubated for 24 h at 37 °C. Centrifugation (3000 × g, 15 min) was done to harvest the bacterial cells. A solution of 0.1% (w/v) phosphate-buffered saline (PBS, Becton Dickinson, New Jersey, US) was used to wash the cell pellets and re-suspended in 50 mL of PBS. For determining the original cell population densities, appropriate dilutions of each cell suspension was made in 0.1% peptone water (PW) and plated in duplicate using Tryptic soy agar (Oxoid Ltd., Basingstoke, UK) plates for *E. coli* suspensions and incubation was done at 37 °C for 24 h. *L. monocytogenes* suspensions were plated on Listeria selective agar base (SR0141E) (Oxoid Ltd., Basingstoke, UK) plates with incubation at 37 °C for 48 h.

### Apple juice inoculation

Aliquots of 45 mL of apple juice were inoculated individually with each of the two bacterial cultures (*L. monocytogenes, E. coli*) targeting a concentration of 10^7^ CFU/mL. The inoculated apple juice was plated using decimal dilutions on Tryptic soy agar (Oxoid Ltd., Basingstoke, UK) plates to determine the original *E. coli* titers and incubation was done at 37 °C for 24 h. Apple juice inoculated with *L. monocytogenes* was plated on Listeria selective agar base (Oxoid Ltd., Basingstoke, UK) and plates were then incubated at 37 °C for 48 h.

### Optical properties and pH measurements

The absorption coefficient at 263 nm was determined based on transmittance measurements from a Cary 300 spectrophotometer with a six-inch integrating sphere (Agilent Technologies, CA, US). Baseline corrections, i.e., by zeroing (setting the full-scale reading of) the instrument using the blank and then blocking the beam with a black rectangular slide was carried out. All pH readings were measured using a standard pH meter (Jenway, Staffordshire, UK). All readings were taken in triplicate to lessen the measurement error. From this data, ultraviolet transmittance value was quantified.

### UV-C irradiation treatments

A collimated beam system that houses a light emitting diode device (Irtronix, Torrence, CA, USA) producing irradiation at 263 nm was used for the exposures. To increase mixing, a 5-mL sample was stirred in a10-mL beakers (height: 3.2 cm; diameter: 1.83 cm) (Bolton and Linden, 2003; Chandra et al., 2017). All optical parameters and UV dose calculations are described in our published studies (Stanley et al., 2020; Islam et al., 2016a; Islam et al., 2016b). Using a high sensitivity sensor (QE Pro series, Ocean Optics, Dunedin, FL, USA), the central irradiance of the UV-C LED system on the surface of the test solution was measured. Based on the central irradiance of the lamp and optical properties (absorption and reduced scattering coefficient) of apple juice, the average fluence was calculated. Four correction factors i.e. reflection (RF) factor, petri factor (PF), divergence factor (DF) and water factor (WF) were accounted in the fluence calculations. UV-C dose was calculated as a product of average fluence and exposure time, equation 2. Correction factors and parameters for obtaining the average fluence rate are shown in Table 2.

**Table 2.**
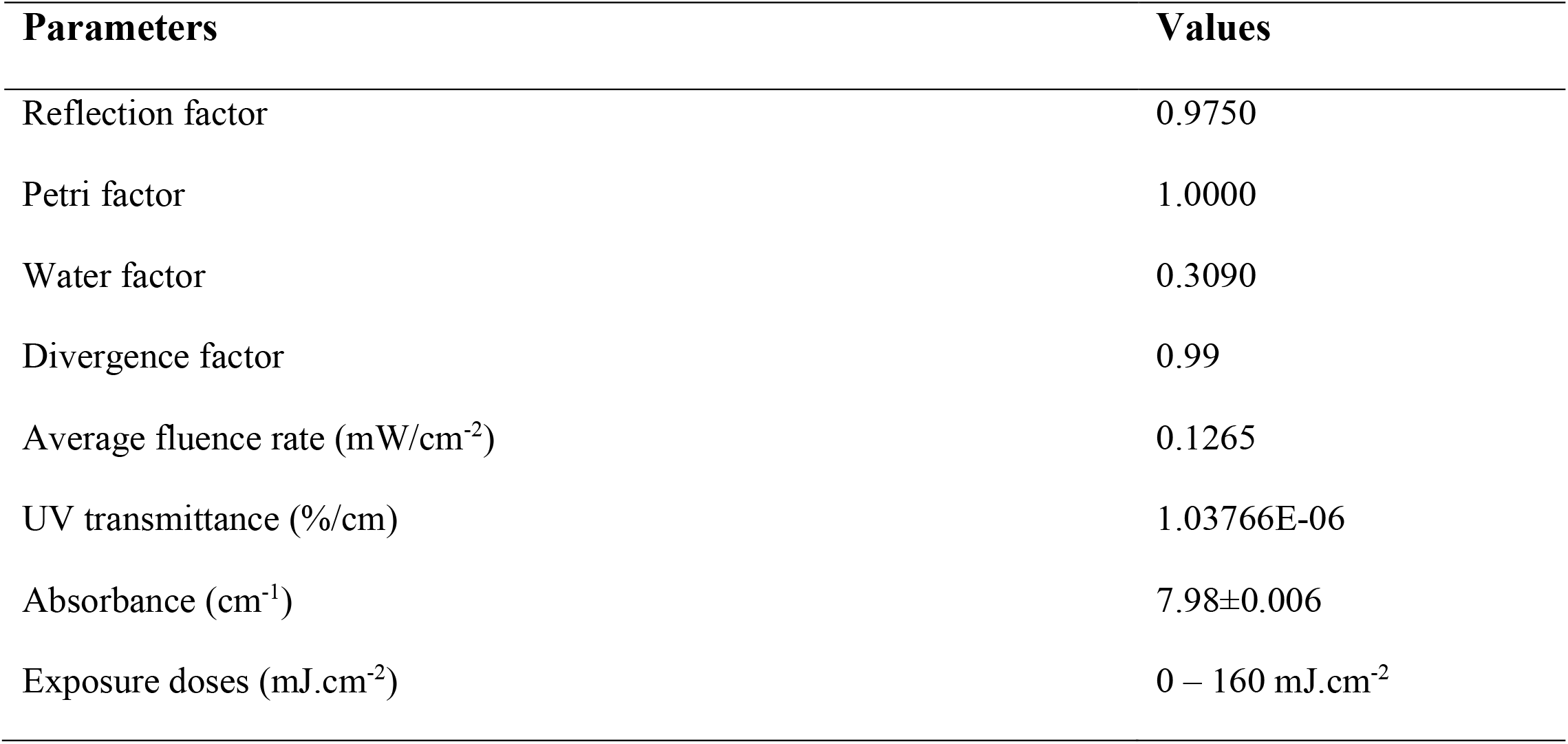
Correction factors and parameters for obtaining the average fluence rate.

The average UV fluence rate in the stirred sample can be calculated as (Eqn. 1):

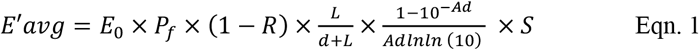

E_0_ is the radiometer meter reading at the center of the beaker and at a vertical position so that the calibration plane of the detector.

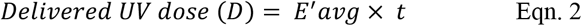

### LC-MS Methods for Polyphenol Detection

A Shimadzu LCMS 8040 system (Shimadzu Scientific Instruments, Columbia, MD) which included two Shimadzu LC-20ADXR pumps, a SIL-20ACXR autosampler, a CTO-20A column oven, and an LCMS-8040 triple stage quadrupole mass spectrometer was used for LC-MS/MS analysis. Chromatographic separation was achieved on with a Phenomenex 1.6 µm Polar C18 100 Å column (100 × 2.1 mm) maintained at temperature of 40°C. The mobile phase consisted of 0.1% formic acid (FA) in water (A) and 0.1% FA in acetonitrile (B). The flow rate was 0.30 mL/min. Initially, solvent B concentration was 3% and increased linearly to 97% from 0.01 min to 2.0 min. Solvent B was held at 97% from 2.0 to 7.0 min, and then reduced to 3% until the end of the time program at 10.0 minutes. The injection volume was 2 µL. The LCMS analysis utilized an electrospray ionization source with the optimized source parameters DL temperature 250 °C, nebulizing gas flow, 3 L/min, heat block 450°C, drying gas flow, 20 L/min. The following transitions and collision energies (CE) were used to analyze compounds in the positive ion mode: pyridoxal m/z 168>150, CE -13; pyridoxamine m/z 168>152, CE -14; pyridoxine m/z 170>152, CE -15; thiamide m/z 264.9>122.05, CE -15; choline m/z 104.1>60, CE-22, Vitamin B12 m/z 678.2>147, CE -40, and chlorogenic acid m/z 354.9>163, CE -15. These compounds were analyzed in the negative ion mode, ascorbic acid m/z 175>114.85, CE 13; phlorizin m/z 535>273.2, CE 15; and epicatechin m/z 289>125, CE 20. A standard solution of 1000 ng/mL each compound was prepared calibration curves. A one-point calibration curve was used to determine concentrations of samples. Data were acquired and analyzed with Shimadzu LabSolutions software.

### Statistics

The concentration of polyphenols, vitamins including microbial inactivation were evaluated under five UV doses levels. A balanced experimental design with three replicates were randomly assigned to each treatment, i.e. exposure to the selected UV-C irradiation. One-way ANOVA test (Tukey’s HSD multiple comparison) was chosen to analyze the data to evaluate the effects of different UV doses on the concentration of polyphenols and vitamins using SAS computing environment. Data were reported as means ± one standard deviation from the mean and tests were statistically significant at 5% significance level.

## Results and Discussion

From the optical data (Table 2), it may be seen that the apple juice was a strong absorber (7.98 cm^- 1^) of UV-C light. On the contrary, some fluids have absorbance between value 0.1 to 22 (Patras et al., 2020; Pendyala et al., 2021; Vashisht et al., 2022). Figure 1 illustrates the ultraviolet (UV) Spectra of apple juice at 254, 263, and 279 nm wavelengths. It clearly shows the absorbance at 263 and 279 nm is relatively lower in comparison to 254 nm wavelength. For beverage treatment, there are some specific system requirements for the use of UV-C light. The near collimated beam (CB) test helps in delivering accurate UV-C dose/fluence for water and wastewater, and it has been considered a standard method (Kuo et al., 2003; Qualls et al., 1983). It is well documented that UV-C exposure is notably altered by optical attenuation coefficient of the test fluid. Perhaps, the dose should be uniformly delivered to the test fluid to reach considerable log reductions of bacteria (Islam et al., 2016b; Patras et al., 2020). This clearly demonstrates that the log reduction obtained data at different absorbances can be considered for comparison only if absorbance is considered and the delivered UV doses were appropriately calculated (Caminiti, et al., 2012; Unluturk et al., 2010, Assatarakul et al., 2012). In this study, all UV gradients in the system were accounted for in the UV dose calculations.

**Figure 1.**
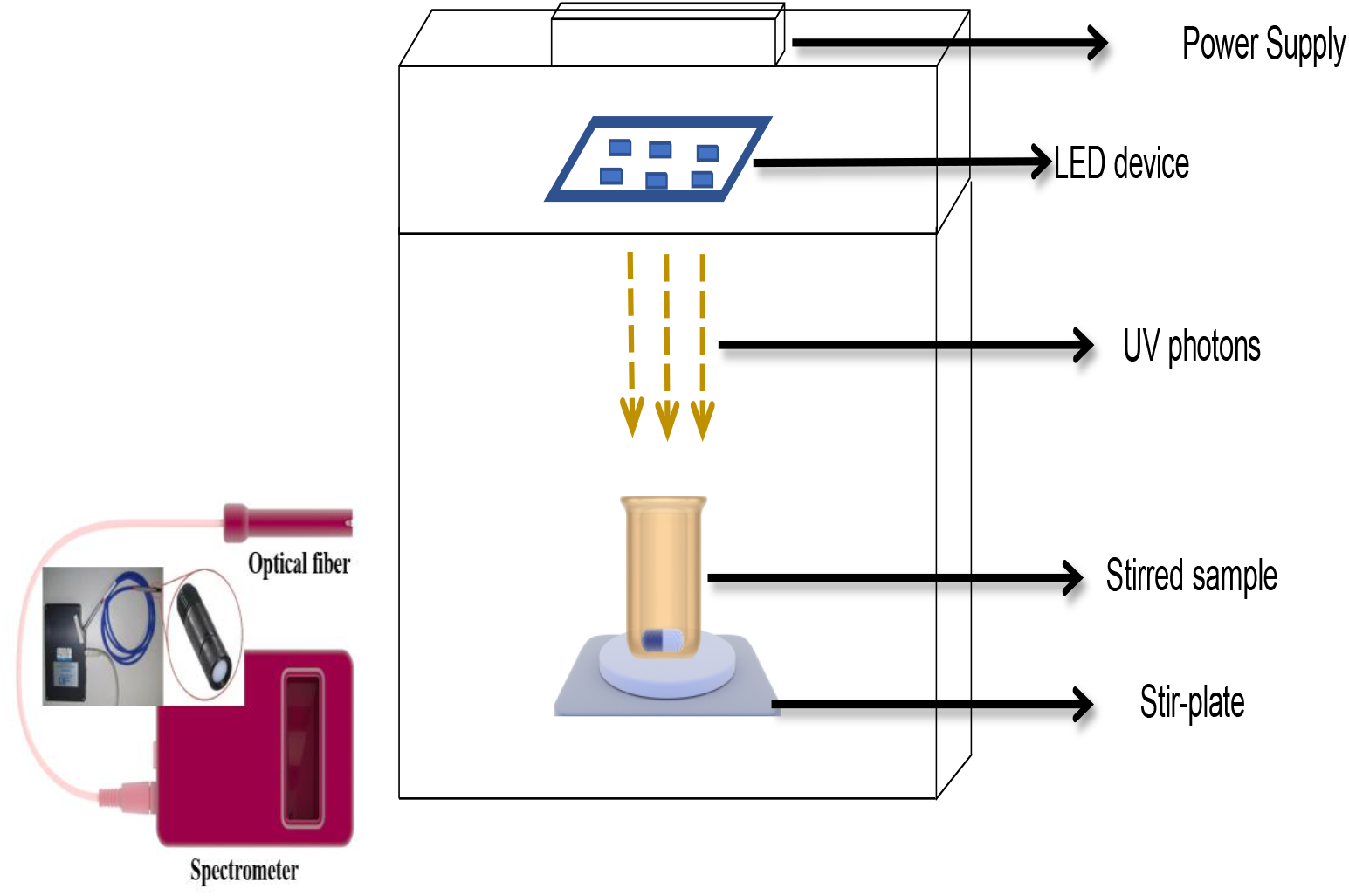
Collimated Light Emitting Diode UV system operating at 263 nm wave-length

**Figure 2.**
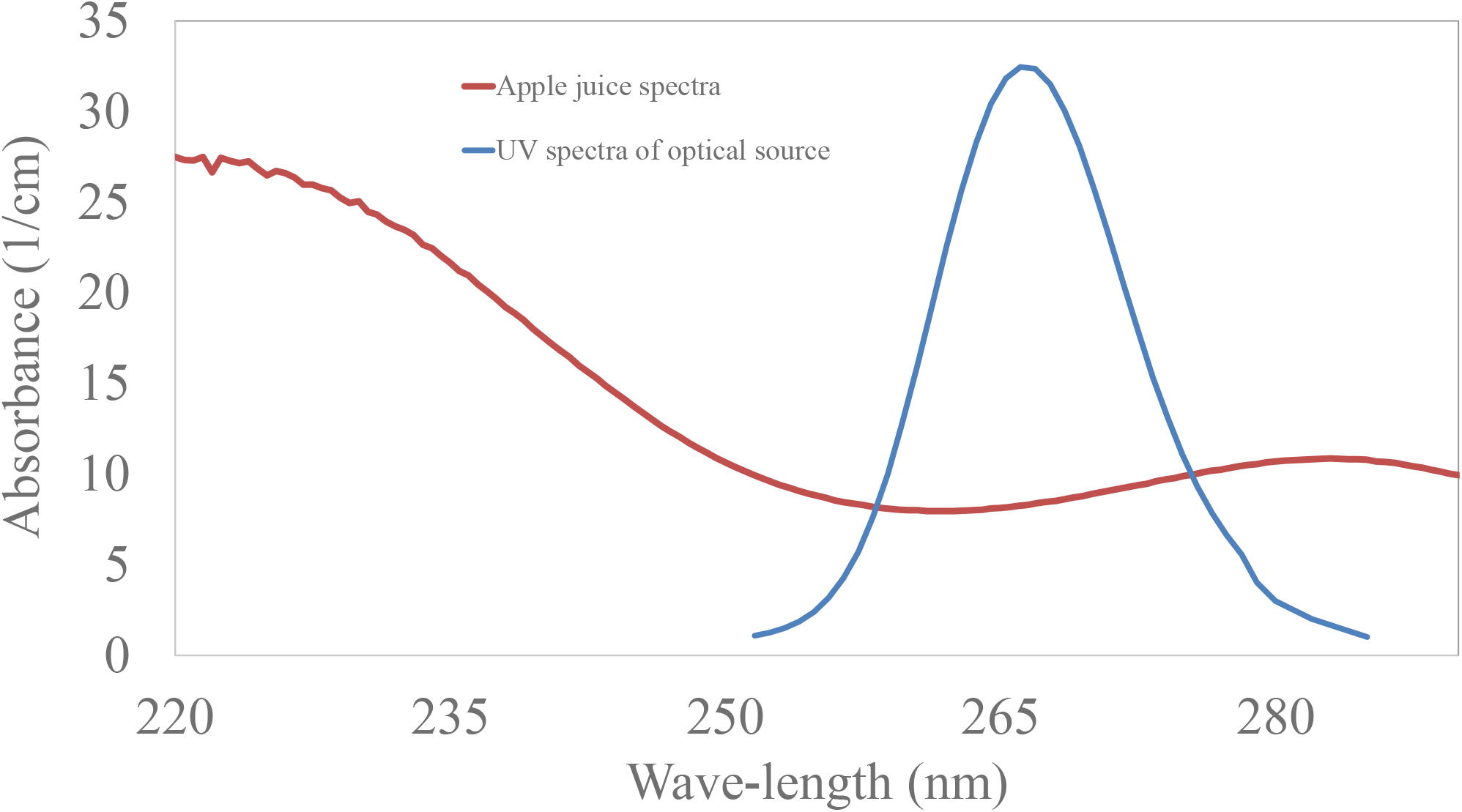
UV Absorbance full scan spectra of apple Juice between 220 – 300nm. UV intensity spectra of Light Emitting Diode UV system, triplicate UV scans were performed all replicates shown on plot

The UV-C inactivation of various pathogens has been studies extensively by many authors (Tosa & Hirata, 1999; Yaun et al., 2003; Sommer et al., 2000; Wilson et al., 1992; Wu et al., 2011). For example, Wilson et al. (1992) observed that *E. coli* O157:H7 ATCC 43894 had an initial shoulder with slow inactivation (initial reduction), followed by first-order kinetics up to 5-log_10_, with D_10_ in the log-linear region of about 1.2 mJ·cm^-2^. It is also worth mentioning that most microbial challenge studies are carried out at 254 nm (Vashisht et al., 2022; Sauceda-Gálvez et al., 2021; Bhullar et al., 2019; Usaga et al., 2017; Chandra et al., 2017; Gunter-Ward et al., 2017; Torkamani, 2011), limited data is available at 263 nm wavelength. Akgün and Ünlütürk, 2017 studied the effects of Ultraviolet light-emitting diodes (UV-LEDs) on the inactivation of *E. coli* K12 (ATCC 25253), an indicator organism of *E. coli* O157:H7 in cloudy apple juice. The clear (AJ) and cloudy apple juice were exposed to UV rays for 40 min by using a UV device composed of four UV-LEDs with peak emissions at 254 and 280 nm and coupled emissions as follows: 254/365, 254/405, 280/365, 280/405 and 254/280/365/405 nm. The authors observed that highest inactivation *of E. coli* K12 (2.0 ± 0.1 log10 CFU/mL and 2.0 ± 0.4 log_10_ CFU/mL) was achieved when the cloudy apple juice was treated with both 280 nm and 280/365 nm UV-LEDs. Perhaps the authors did not quantify the delivered dose and did not validate the dosage. In a different study, Chevremont et al. (2012a) showed that coupling UV-A and UV-C could be paired by using the germicidal effect of UV-C and greater penetrating ability of UV-A. They also found that use of coupled wavelengths 280/365 nm and 280/ 405 nm caused total disappearance of fecal enterococci, total coliforms and fecal coliforms in the effluent. Besides lack of possibility to repair the damage in the bacterial membranes occurred after UV-A exposure increased the efficiency of microbial inactivation (Chevremont et al., 2012a). Ngadi et al. (2003) applied 300 mJ/cm^2^ UV fluence at 0.32 mW/cm^2^ incident UV intensity for 16 min in order to achieve 4.2 log_10_ (CFU/mL) reduction of *E. coli* O157:H7 (ATCC 35150) in apple juice samples of 0.1 cm in optical depth. The authors irradiated the samples using a collimated beam apparatus consisted of a low-pressure mercury UV lamp with peak radiation in the 255.7 nm wavelength range. In a separate study, apple juice (absorption coefficient 5.81 cm^-1^) inoculated with *E. coli* K12 was treated with UV-C irradiation and 4.6 log_10_ CFU/mL reduction was obtained (Caminiti et al., 2012). In our study, 10 mJ/cm^2^ UV dose at 0.288645 mW/cm^2^ average irradiance resulted in 3.4 log_10_ CFU/mL reduction of *E. coli* at a depth of 1.5 cm using UV-LEDs emitting light at 263 nm. Fluid absorbance is an important parameter that affects the UV light intensity and its penetrating ability. This study rectifies this issue and accounts for UV intensity gradients in the dose calculations. The current study examines the inactivation of *Escherichia coli* and *Listeria monocytogenes* in natural apple juice, for which the authors found limited published literature regarding 263nm wavelength or closer exposure wavelengths.

The populations of *Escherichia coli* were reduced by 1.03, 1.87, 2.49, 3.45 log_10_ respectively at a UV-C dose level of 0, 4, 8, 10, 15 mJ·cm^-2^. The microbial inactivation shows first order kinetics, as plotted in Fig. 3. Log inactivation was proportional to UV dosage. Based on the rate constant (cm^2^/mJ) and D_10_ value of *Escherichia coli*, 20.8 mJ.cm^-2^ dosage will be required to achieve 5 log_10_ reduction. In our bench scale experiments, apple juice samples were continuously stirred during the entire duration of the irradiation treatments to ensure uniform dose delivery to a homogeneous test fluid. Log inactivation increased with increase in UV-C exposure, as evidenced in Fig 3. This may be foreseen as the fact that UV-C light inactivates microbial population by preventing replication of DNA and cellular damage. In some cases, if the dose is not delivered uniformly, UV-C dose beyond the threshold capacity of cellular damage results in rapid lethal inactivation of cells and DNA repair mechanisms fail to undo changes (Miller et al., 1999). Previous studies reported D values between 1.5 – 6 mJ/cm^2^ for 1 and 4 log_10_ reductions (Tosa and Hirata, 1999); and <2 mJ/cm^2^ through 17 mJ/cm^2^ respectively representing between 1 and 6 total log reduction (Yaun et al., 2003) for *Escherichia coli O157:H7, and for Escherichia coli* O157:H7 ATCC 43894 the d values ranged between 1.5 and 6.8; which is equivalent to between a one and five-log reduction (Wilson et al., 1992). All data is in accordance with the published studies except high d values. The higher d values could be due to poor mixing in the system due to false tailing effects. In the conventional approach of using near collimated beam apparatus for UV-C exposure, it is assumed that the mixing is adequate for all micro-organisms to accumulate the same UV-C dose. One of the assumptions used in the calculation of UV-C dose-response data is that all of the fluid elements receive the same dose under vigorous mixing. This assumption may fail in some scenarios where fluid and micro-organisms are different.

**Figure 3:**
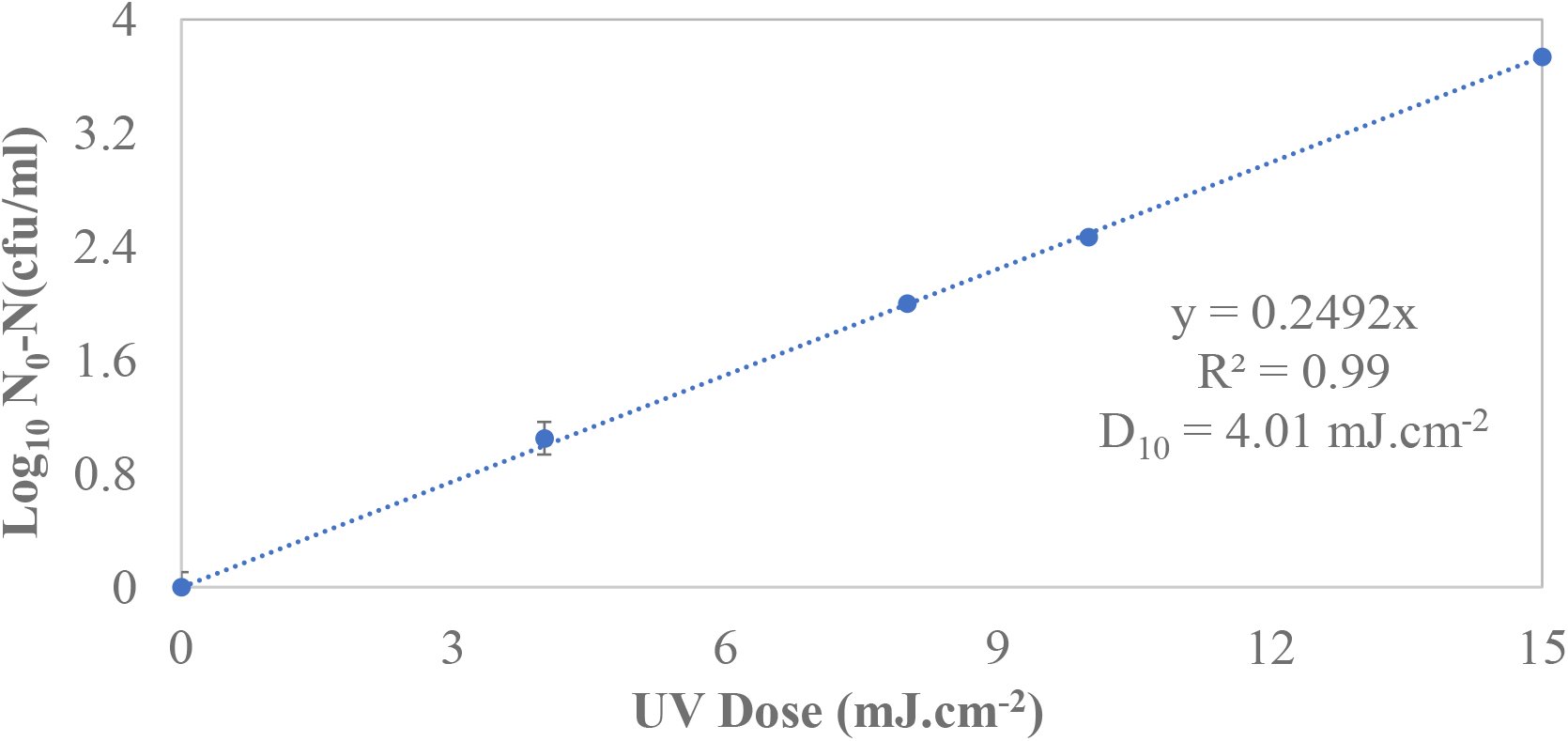
UV-C Inactivation of *Escherichia coli* O157:H7 in apple juice using a collimated Light Emitting Diode UV system at 263 nm wave-length; the fluence intensity gradients were adjusted (i.e. optical properties of apple juice). Triplicate irradiations were performed for each dose; all replicates shown on plot, and values shown are averages of duplicate plating of each irradiated sample. Error bars represent range of data.

UV-C irradiation effectively inactivated *L. monocytogenes* ATCC 19115 in apple juice as may be seen in Fig. 4. The populations of *L. monocytogenes* were reduced by 1.25, 2.37, 3.26, 4.13 log_10_ respectively at a UV-C dose level of 4, 8, 12, 15 mJ·cm^-2^. The D_10_ value obtained overall in the apple juice was 3.60 mJ/cm^-2^. The graph demonstrates that when the UV dosage is increased, the log reduction is increasing. Based on the rate constant (cm^2^/mJ) and D_10_ value of *L. monocytogenes*, 18.12 mJ.cm-^2^ dosage will be required to achieve 5 log_10_ reduction. Literature data suggests that *L. monocytogenes* UV sensitivity lies between 3.24 mJ.cm^-2^ (Gunter Ward et al., 2017), 4 mJ.cm^-2^ log reduction (Lu et al., 2010), and greater than 5 mJ.cm^-2^ (Matak et al., 2005). Higher d value could be due to poor mixing in the system, creating huge UV gradients, giving a non-uniform dose distribution. At the maximum regulated UV-C dose set by the FDA (40 mJ·cm^-2^), the predicted inactivation treatments can result in a 5-log_10_ decrease of E. *coli*, and *L. monocytogenes* in apple juice. To conclude, this system almost allowed accurate estimation of the delivered doses required to inactivate 5 log_10_ reductions of these pathogens taking into account the UV-C irradiance, absorption coefficient (1/cm). Kinetic data shows that both pathogens are very sensitive to UV light at 263nm wavelength. Overall, in this study, 20 mJ/cm^-2^ of exposure treatment is the optimized dosage for safer and higher quality product.

**Figure 4:**
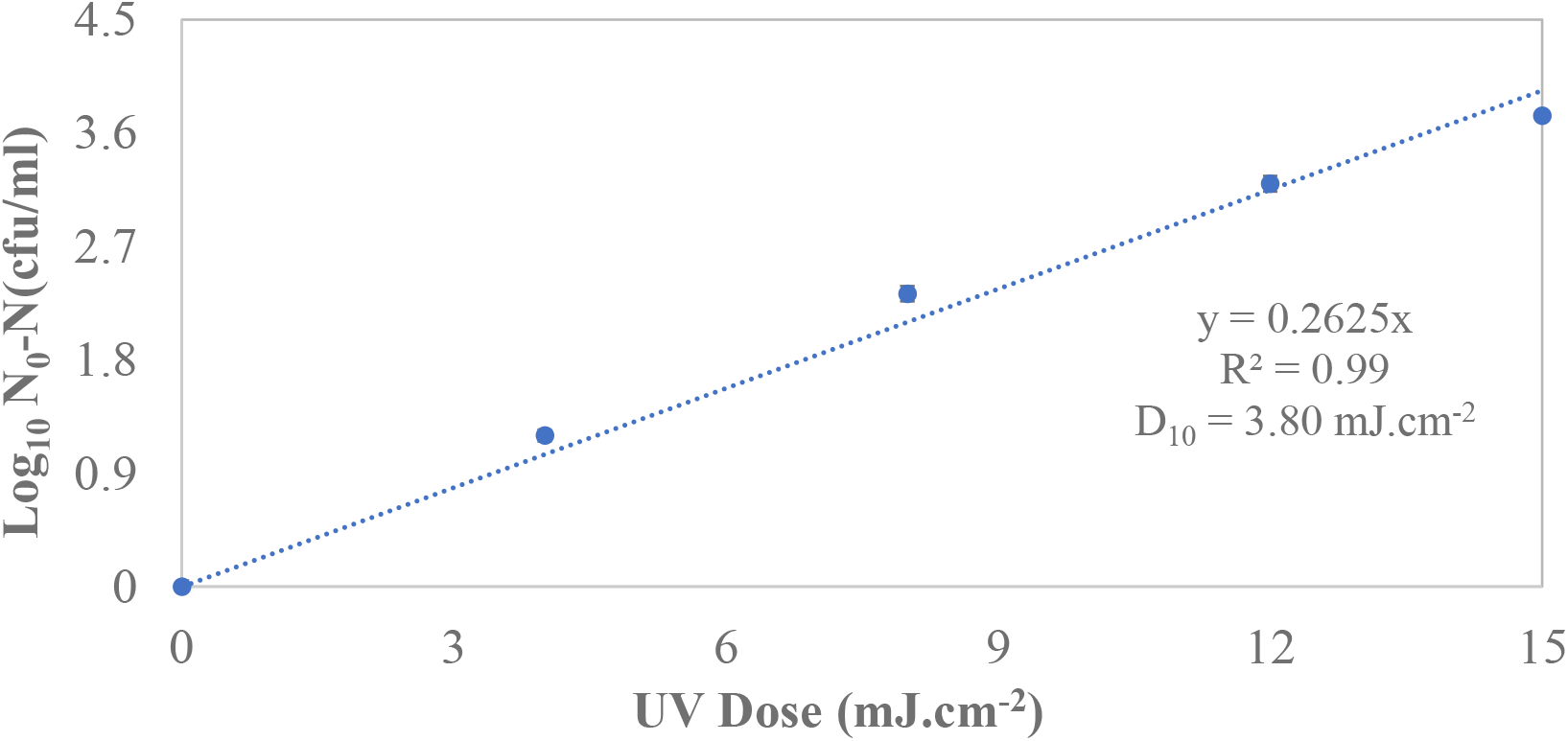
UV-C Inactivation of and *Listeria monocytogenes* in apple juice using a collimated Light Emitting Diode UV system at 263 nm wave-length; the fluence intensity gradients were adjusted (i.e. optical properties of apple juice). Triplicate irradiations were performed for each dose; all replicates shown on plot, and values shown are averages of duplicate plating of each irradiated sample. Error bars represent range of data.

Figures 5 and 6 illustrate the effect of UV-C irradiation on the content of polyphenolic and vitamin compounds present in apple juice. Figure 7 demonstrate chromatograms of ascorbic acid and epicatechin respectively Chlorogenic acid has notably demonstrated as being the most abundant of polyphenolic compounds present in apple juice (Islam et al., 2016b; Eisele and Drake 2005; Islam et al., 2016a). Another major polyphenol in apple juice is epicatechin (Marcotte, et al., 2022; Boyer and Liu, 2004). Concentration of epicatechin was observed to diminish as a function of UV dosage (p < 0.05). At the maximum dosage level (160 mJ.cm^-2^), epicatechin and chlorogenic acid were significantly reduced by 98% and 76% respectively. Results suggested that epicatechin was relatively sensitive to UV-C dosage as compared to chlorogenic. Perhaps, approximately, 30% reduction was observed at 20 mJ.cm^-2^ for both polyphenols.

**Figure 5.**
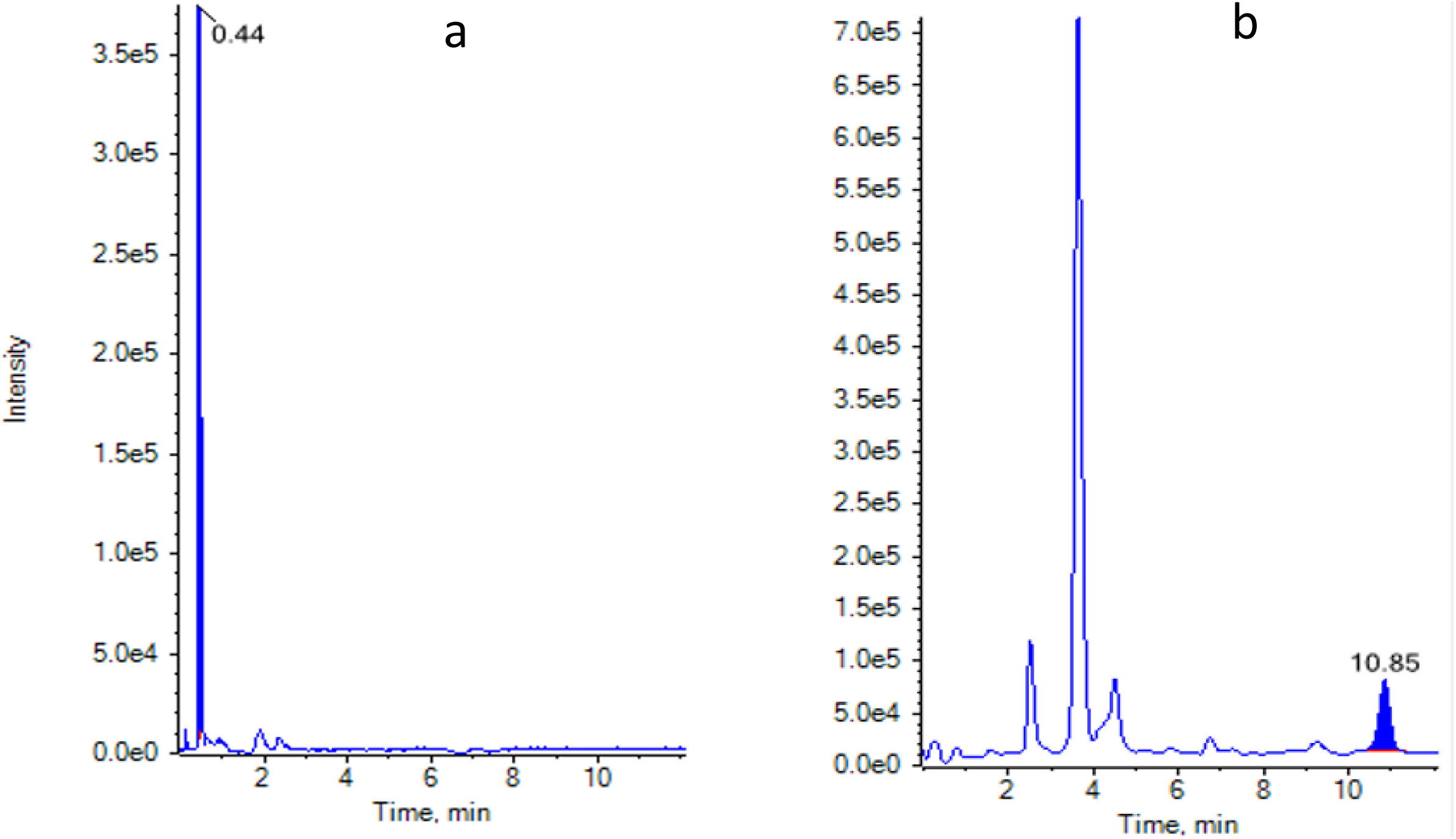
Representative Chromatograms of Ascorbic Acid (a) and Epicatechin (b).

**Figure 6.**
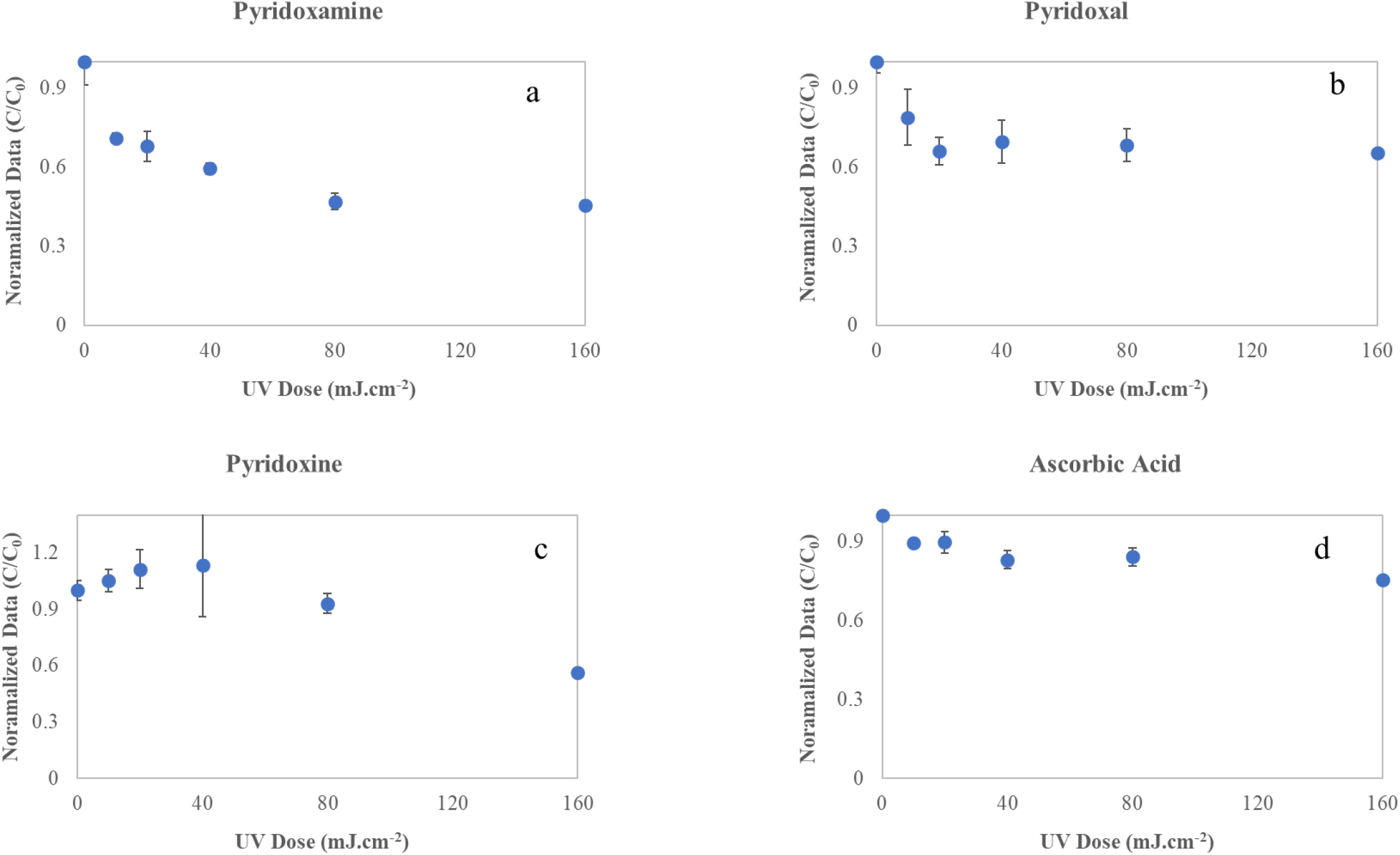
Effect of UV irradiation on the stability of vitamins [a. Pyridoxamine; b. Pyridoxal; c. Pyridoxine; d. Ascorbic acid] as a function of UV dose. The plots contain aggregated data from multiple experiments in which exposures were performed three times at each level. Error bars represent range of data

**Figure 7.**
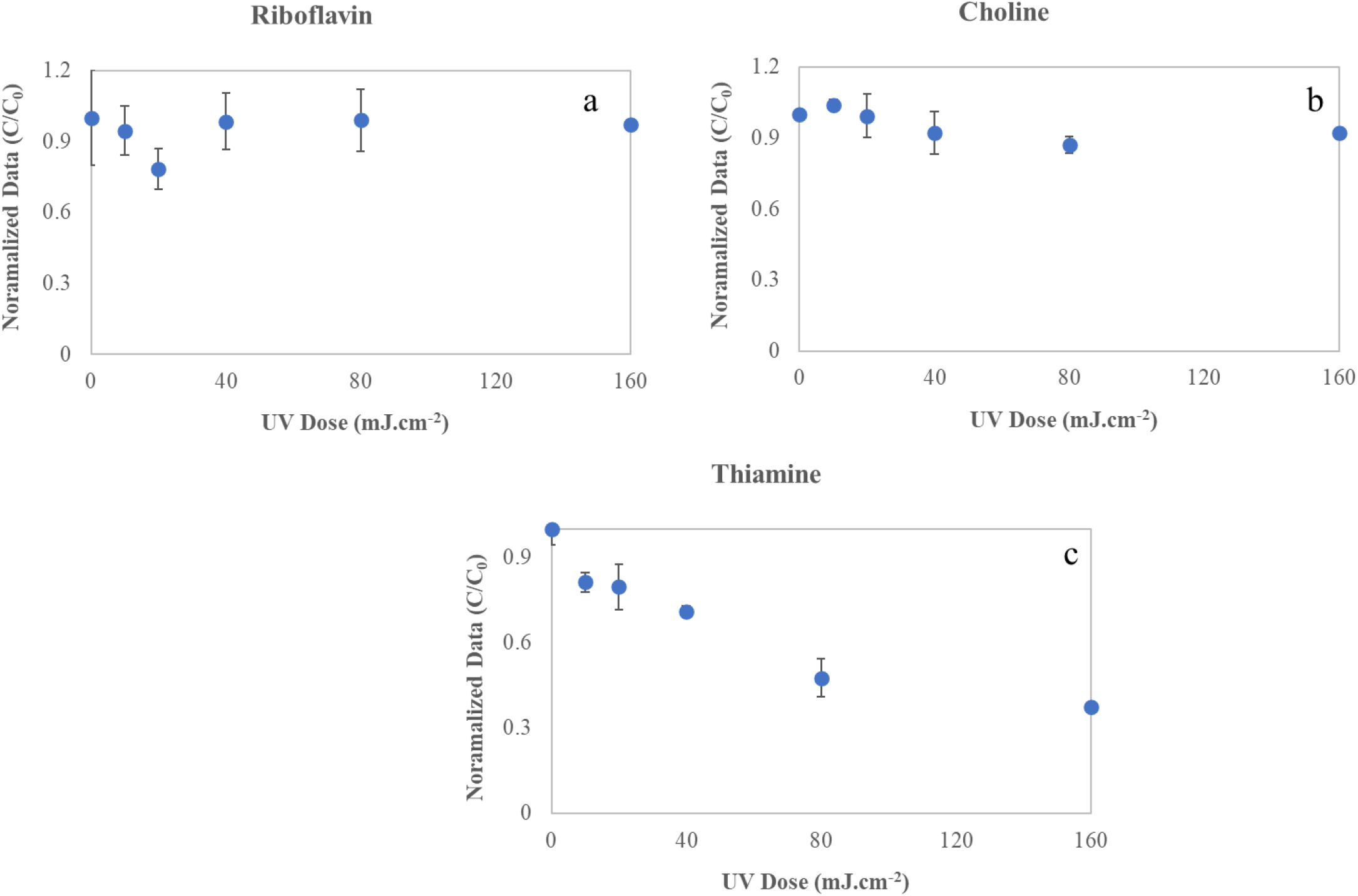
Effect of UV irradiation on the stability of vitamins [a. Riboflavin; b. Choline; c. Thiamine]. as a function of UV dose. The plots contain aggregated data from multiple experiments in which exposures were performed three times at each level. Error bars represent range of data

**Figure 8.**
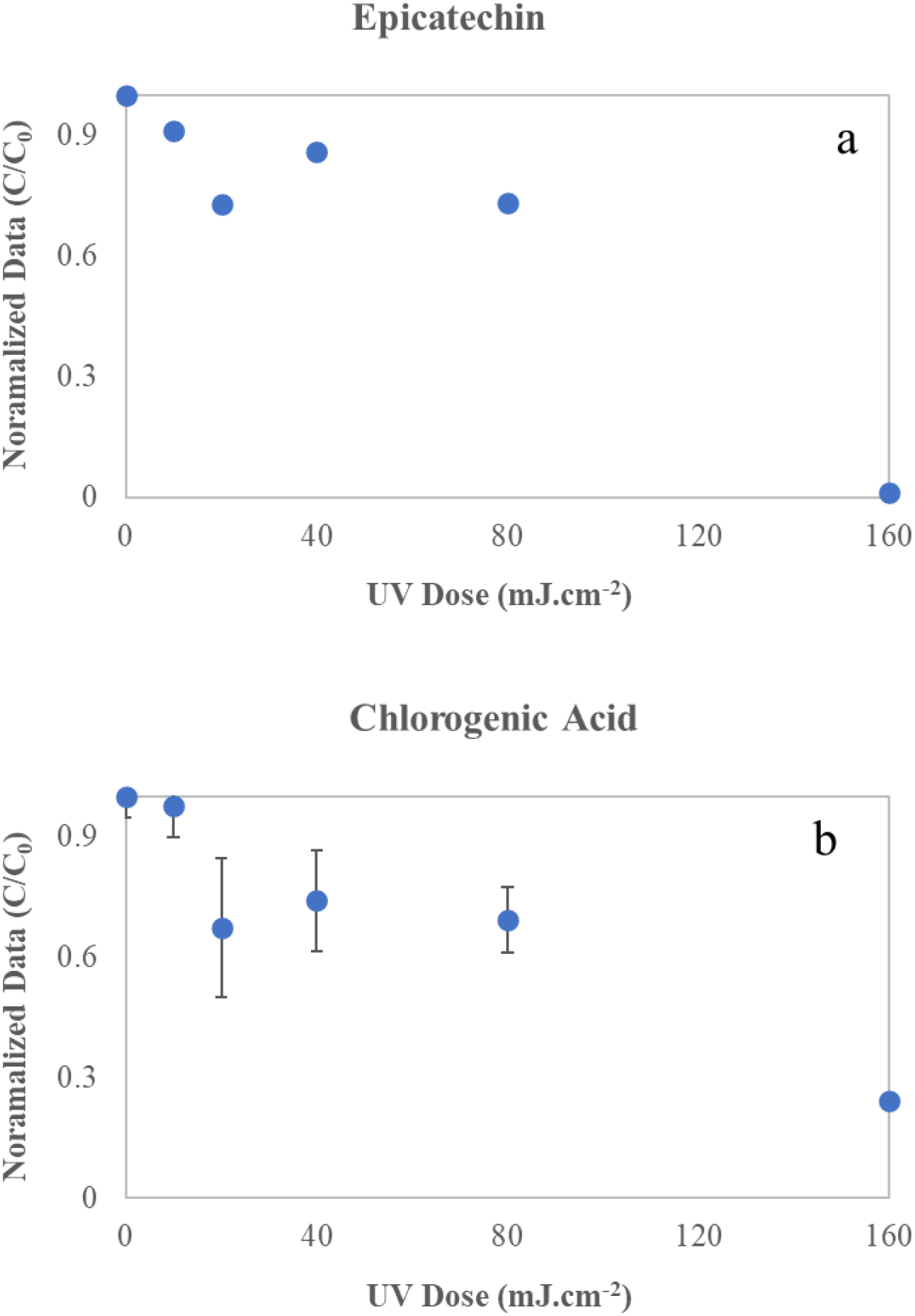
Effect of UV irradiation on the stability of polyphenols [a. Epicatechin; b. Chlorogenic acid] as a function of UV dose. The plots contain aggregated data from multiple experiments in which exposures were performed three times at each level. Error bars represent range of data

Seven vitamins were identified and quantified in apple juice by LC-MS/MS; all of the vitamins identified were significantly reduced (p < 0.05) as UV dosage was increased. Pyridoxamine (B6), pyridoxal (B6), pyridoxine (B6), chlorogenic acid, riboflavin (B2), Epicatechin, Thiamine (B1), Choline (B7). Riboflavin (Min et al., 2002a; Min et al., 2002b), Thiamine (Whitfield et al., 2018), and Pyridoxine (Kawada et al., 2000) are all very photosensitive vitamins and act as photosensitizers due to the presence of the double bonds located within their structures, while also accelerating the oxidation of other B vitamins (De Arrivetti et al., 2013). Riboflavin is a vitamin that is very photosensitive with the capability to act as a photosensitizer because of the double bonds housed within its structure, (Min et al., 2002; Min and Boff, 2002). it can accelerate the oxidation of other B vitamins (De O.R. Arrivetti et al., 2013). In this study, riboflavin concentrations in the samples decreased by 5.40, 21.6, 1.57, 0.094, and 0.029% at UV doses at 10, 20, 40, 80 and 160 mJ.cm^-2^, respectively. Riboflavin is rapidly degraded by UV-C exposures. Similar findings were reported in another study of UV light on Nutritional Quality and Cytotoxicity of Apple Juice in (Islam et al., 2016 a), UV-C irradiation as an alternative disinfection method and the effect of UV-C in polyphenols and apple juice antioxidant activity in (Islam et al., 2016 b). Based on concentration reductions, it appears that all vitamins were very sensitive to UV-C exposures in apple juice and decreased significantly. The degradation of vitamins during UV irradiation may result from a various possible chemical reactions and mechanisms as noted in Islam et al., 2016. Degradation of vitamins may be related to the photo-oxidation when oxygen and other factors such as light are present (Islam et al, et al., 2017). In contrast, ascorbic acid was retained well in UV exposed samples, whereas the concentration of thiamine was reduced significantly. Our study demonstrated that UV-C irradiation caused significant changes in vitamins and polyphenols, and it is quite indicative that UV irradiation is capable to change a number of chemical food constituents. To the best of the authors’ knowledge, this is the first paper that demonstrates the efficacy of UVC 263 nm inactivating *Escherichia coli*, and *Listeria monocytogenes* in highly turbid apple juice and retaining the quality between 20-40 mJ.cm^-2^ dosage.

## Conclusion

UV-C light reduced the microbial load in apple juice, making it a promising alternative pasteurization technique. UV-C fluence of 12 mJ·cm^-2^ achieved 3.5 -log reduction (99.9%) in *Escherichia coli* ATCC 700728 and *Listeria monocytogenes* respectively. The D_10_ values of 4.16 and 3.84 mJ·cm-2 were obtained for *Escherichia coli*, and *Listeria monocytogenes*. The log reduction kinetics of microorganisms followed log-linear and with higher R^2^ (>0.95) and low RMSE values. UV-C irradiation induced minor reductions in the concentration of vitamins and polyphenols in apple juice at the FDA-recommended fluence of 20-40 mJ cm^-2^ of pasteurized equivalent dose. Ascorbic acid was reduced to 32%, at 160 mJ/cm^2^ whereas 17% reduction was observed at 40 mJ/cm^2^ (quality indicator). Riboflavin was observed to be relatively stable as a function of UV dosage. Although our research findings are not specifically suitable to all juices, due to complex photochemistry of different fluid matrices.

## Acknowledgments

The authors would like to thank Mrs. Yvonne Myles, Dr. Jerwzy, Mr. Vybhav Gopisetty, and Ms. Judy Stanley for providing valuable guidance in this project.

## Credit Authorship Contribution Statement

Brahmaiah Pendyala: Conceptualization, Methodology, and Investigation.

Ankit Patras: Conceptualization, Methodology, Supervision, All draft reviews, Funding acquisition. Fur-Chi Chen: Conceptualization, Methodology, Supervision, and Investigation. Anjali Kurup: Conceptualization, Methodology, and Investigation.

Matthew Vergne: Conceptualization, Methodology, and Investigation.

Vybhav Vipul Sudhir Gopisetty: Methodology, and Conceptualization. Judy Stanley: Methodology, and Conceptualization. Yvonne Myles: Conceptualization, Methodology, and Investigation. Anita Scales Akwu: Methodology, Investigation.

## References

Akgün, M. P., & Ünlütürk, S. (2017). Effects of ultraviolet light emitting diodes (LEDs) on microbial and enzyme inactivation of apple juice. International Journal of Food Microbiology, 260, 65–74.

Assatarakul, K., Churey, J. J., Manns, D. C., & Worobo, R. W. (2012). Patulin reduction in apple juice from concentrate by UV radiation and comparison of kinetic degradation models between apple juice and apple cider. Journal of Food Protection, 75(4), 717–724.

Bao, M.-J., Shen, J., Jia, Y.-L., Li, F.-F., Ma, W.-J., Shen, H.-J., Shen, L.-L., Lin, X.-X., Zhang, L.-H., Dong, X.-W., Xie, Y.-C., Zhao, Y.-Q., & Xie, Q.-M. (2013). Apple polyphenol protects against cigarette smoke-induced acute lung injury. Nutrition Journal, 29(1), 235–243.

Bhullar, M. S., Patras, A., Kilonzo-Nthenge, A., Pokharel, B., & Sasges, M. (2019). Ultraviolet inactivation of bacteria and model viruses in coconut water using a collimated beam system. Food Science and Technology International, 25(7), 562–572.

Bolton, J. R., & Linden, K. G. (2003). Standardization of Methods for Fluence (UV Dose) Determination in Bench-Scale UV Experiments. Journal of Environmental Engineering, 129(3), 209–216.

Boyer, J., & Liu, R. H. (2004). Apple phytochemicals and their health benefits. Nutrition Journal, 3(5).

Caminiti, I. M., Palgan, I., Muňoz, A., Noci, F., Whyte, P., Morgan, D. J., Cronin, D. A., & Lyng, J. G. (2012). The effect of ultraviolet light on microbial inactivation and quality attributes of apple juice. Food Bioprocess Technology, 5.

(CFSAN), C. F. S. A. N. (2000). Kinetics of Microbial Inactivation for Alternative Food Processing Technologies. (IFT/FDA Contract No. 223-98-2333). United States Food and Drug Administration, United States Department of Health and Human Services Retrieved from https://www.fda.gov/files/food/published/Kinetics-of-Microbial-Inactivation-for-Alternative-Food-Processing-Technologies.pdf.

Chandra, S., Patras, A., Pokharel, B., Bansode, R. R., Begum, A., & Sasges, M. (2017). Patulin degradation and cytotoxicity evaluation of UV irradiated apple juice using human peripheral blood mononuclear cells. Journal of Food Process Engineering, 40(6).

Chevremont, A. C., Farnet, A. M., Coulomb, B., & Boudenne, J. L. (2012). Effect of Coupled UV-A and UV-C LEDs on both Microbiological and Chemical Pollution of Urban Wastewaters. Science of the Total Environment, 426, 304–310.

De Arrivetti, O. R. L., Scurachio, R. S., Santos, W. G., Papa, T. B. R., Skibsted, L. H., & Cardoso, D. R. (2013). Photooxidation of Other B Vitamins as Sensitized by Riboflavin. Journal of Agricultural and Food Chemistry, 61(31), 7615–7620.

Eisele, T. A., & Drake, S. (2005). The partial compositional characteristics of apple juice from 175 apple varieties. Journal of Food Composition and Analysis, 18(2& 3), 213–221.

Franz, C. M. A. P., Specht, I., Cho, G. S., Graef, V., & Stahl, M. R. (2009). UV-C-inactivation of microorganisms in naturally cloudy apple juice using novel inactivation equipment based on Dean vortex technology. Food Control, 20 (12), 1103–1107.

Gunter-Ward, D. M., Patras, A., Bhullar, M. S., Kilonzo-Nthenge, A., Pokharel, B., & Sasges, M. (2017). Efficacy of ultraviolet (UV-C) light in reducing foodborne pathogens and model viruses in skim milk. Journal of Food Processing and Preservation, 42(2).

Islam, M. S., Patras, A., Pokharel, B., Wu, Y., Vergne, M. J., Shade, L., Xiao, H., & Sasges, M. (2016a). UV-C irradiation as an alternative disinfection technique: Study of its effect on polyphenols and antioxidant activity of apple juice. Innovative Food Science Emerging Technology, 34, 344–351.

Islam, M. S., Patras, A., Pokharel, B., Vergne, M., Sasges, M., Begum, A., Rakariyatham, K., Pan, C., & Xiao, H. (2016b). Effect of UV Irradiation on the Nutritional Quality and Cytotoxicity of Apple Juice. Journal of Agricultural and Food Chemistry, 64(41), 7812–7822.

Kahle, K., Kraus, M., & Richling, E. (2005). Polyphenol profiles of apple juices. Molecular Nutrition & Food Research, 49(8), 797–806.

Kawada, A., Kashima, A., Shiraishi, H., Gomi, H., Matsuo, I., Yasuda, K., Sasaki, G., Sato, S., & Orimo, H. (2000). Pyridoxine-Induced Photosensitivity and Hypophosphatasia. Dermatology, 201(4), 356–360.

Kaya, Z., Yıldız, S., & Ünlütürk, S. (2015). Effect of UV-C irradiation and heat treatment on the shelf life stability of a lemon-melon juice blend: multivariate statistical approach. Innovative Food Science & Emerging Technologies, 29, 230–239.

Keyser, M., Muller, I. A., Cilliers, F. P., Nel, W., & Gouws, P. A. (2008). Ultraviolet radiation as a non-thermal treatment for the inactivation of microorganisms in fruit juice. Innovative Food Science and Emerging Technologies, 9(3), 348–354.

Khanizadeh, S., Taso, R. V., Rekika, D., Yang, R., Charles, M. T., & Rupasinghe, H. P. V. (2008). Polyphenol composition and total antioxidant capacity of selected apple genotypes for processing. Journal of Food Composition and Analysis, 21(5), 396–401.

Kschonsek, J., Wolfram, T., Stöckl, A., & Böhm, V. (2018). Polyphenolic compounds analysis of old and new apple cultivars and contribution of polyphenolic profile to the in vitro antioxidant capacity. Antioxidants (Basel), 7(1).

Kuo, J., Chen, C.-l., & Nellor, M. (2003). Discussion of “Standardized Collimated Beam Testing Protocol for Water/Wastewater Ultraviolet Disinfection” Journal of Environmental Engineering, 129(8), 774–779.

Kurup, A. H., Patras, A., Pendyala, B., Vergne, M. J., & Bansode, R. R. (2022). Evaluation of Ultraviolet-Light (UV-A) Emitting Diodes Technology on the Reduction of Spiked Aflatoxin B1 and Aflatoxin M1 in Whole Milk. Food and Biopress Technology, 15, 165–176.

Lu, G., Li, C., Liu, P., Cui, H., Yao, Y., & Zhang, Q. (2010). UV inactivation of microorganisms in beer by novel thin-film apparatus. Food Control, 21(10), 1312–1317.

Marcotte, B. V., Verheyde, M., Pomerleau, S., Doyen, A., & Couillard, C. (2022). Health Benefits of Apple Juice Consumption: A Review of Interventional Trials on Humans. Nutrients, 14(4), 821.

Markowski, J., Baron, A., Mieszczakowska, M., & Płocharski, W. (2009). Chemical composition of French and Polish cloudy apple juices. The Journal of Horticultural Science and Biotechnology, 84(6), 68–74.

Matak, K. E., Churey, J. J., Worobo, R. W., Sumner, S. S., Hovingh, E., Hackney, C. R., & Pierson, M. D. (2005). Efficacy of UV-C light for the reduction of Listeria monocytogenes in goat’s milk. Journal of Food Protection, 68, 2212–2216.

Miller, D. N., Bryant, J. E., Madsen, E. L., & Ghiorse, W. C. (1999). Evaluation and optimization of DNA extraction and purification procedures for soil and sediment samples. Applied and Environmental Microbiology, 65(11), 4715–4724.

Min, D. B., Boff, J. M., & Akoh, C.C. (2002). Lipid oxidation of edible oil (2nd ed.). The Ohio State University.

Min, D. B., & Boff, J. M. (2002). Chemistry and reaction of singlet oxygen in foods. Comprehensive Reviews in Food Science and Food Safety, 1, 58–72.

Müller, A., Noack, L., Greiner, R., Stahl, M. R., & Posten, C. (2014). Effect of UV-C and UV-B treatment on polyphenol oxidase activity and shelf life of apple and grape juices. Innovative Food Science and Emerging Technologies, 26, 498–504.

Ngadi, M., Smith, J. P., & Cayouette, B. (2003). Kinetics of ultraviolet light inactivation of Escherichia coli O157:H7 in liquid foods. Journal of the Science of Food and Agriculture, 83(15), 1551–1555.

Oszmianski, J., Wolniak, M., Wojdylo, A., & Wawer, I. (2007). Comparative study of polyphenolic content and antiradical activity of cloudy and clear apple juices. Journal of the Science of Food and Agriculture, 87(4), 573–579.

Patras, A., Ricketts, J., Pendyala, B., & Godwin, S. (2020). UV-C Light Ensuring Safety and Quality of Beverages. 1 - 2. https://www.tnstate.edu/extension/documents/UV%20Light%20Fact%20Sheet%20FSBE-01-2020-1.pdf.

Patras, A., Tiwari, B. K., & Brunton, N. P. (2011). Influence of blanching and low temperature preservation strategies on antioxidant activity and phytochemical content of carrots, green beans and broccoli. LWT – Food Science and Technology, 44(1), 299–306.

Pendyala, B., & Patras, A. (2021). Predicted UV-C Sensitivity of Human and Non-human Vertebrate (+) ssRNA Viruses. bioRxiv: The Preprint Server for Biology. https://www.biorxiv.org/content/10.1101/2021.05.10.443521v1.

Qualls, R. G., Flynn, M., & Johnson, J. D. (1983). The Role of Suspended Particles in Ultraviolet Disinfection. Journal of Water Pollution Control Federation, 55(10), 1280–1285.

Sauceda-Gálvez, J. N., Martinez-Garcia, M., Hernández-Herrero, M. M., Gervilla, R., & Roig-Sagués, A. X. (2021). Short Wave Ultraviolet Light (UV-C) Effectiveness in the Inactivation of Bacterial Spores Inoculated in Turbid Suspensions and in Cloudy Apple Juice. Beverages, 7(1).

Sommer, R., Lhotsky, M., Haider, T., & Cabaj, A. (2000). UV inactivation, liquid-holding recovery, and photoreactivation of Escherichia coli O157 and other pathogenic Escherichia coli strains in water. Journal of Food Protection, 63(8), 1015–1020.

Stanley, J., Patras, A., Pendyala, B., Vergne, M. J., & Bansode, R. R. (2020). Performance of a UV-A LED system for degradation of aflatoxins B1 and M1 in pure water: kinetics and cytotoxicity study. Scientific Reports, 10(1), 1–12.

Torkamani, A. E. (2011). Impact of UV-C light on orange juice quality and shelf life. International Food Research Journal, 18(4).

Tosa, K., & Hirata, T. (1999). Photoreactivation of enterohemorrhagic Escherichia coli following UV disinfection. Water Research, 33(2), 361–366.

Unluturk, S., Atilgan, M. R., Baysal, A. H., & Unluturk, M. (2010). Modeling inactivation kinetics of liquid egg white exposed to UV-C irradiation. International Journal of Food Microbiology, 142, 341–347.

Usaga, J., Manns, D. C., Moraru, C. I., Worobo, R. W., & Padilla-Zakour, O. (2017). Ascorbic acid and selected preservatives influence effectiveness of UV treatment of apple juice. Lebensmittel-Wissenschaft & Technologie (LWT), 75, 9–16.

Whitfield, K. C., Bourassa, M. W., Adamolekun, B., Bergeron, G., Bettendorff, L., Brown, K. H., Cox, L., Fattal-Valevski, A., Fischer, P. R., Frank, E. L., Hiffler, L., Hlaing, L. M., Jefferds, M. E., Kapner, H., Kounnavong, S., Mousavi, M. P. S., Roth, D. E., Tsaloglou, M.-N., Wieringa, F., & Combs, J. G.F.. (2018). Thiamine deficiency disorders: diagnosis, prevalence, and a roadmap for global control programs. Annals of the New York Academy of Sciences, 1430(1), 3–43.

Van der Sluis, A. A., Dekker, M., Skrede, G., & Jongen, W. M. F. (2002). Activity and concentration of polyphenolic antioxidants in apple juice effect of existing methods. Journal of Agricultural and Food Chemistry, 50(25), 7211–7219.

Vashisht, P., Pendyala, B., Patras, A., Gopisetty, V. V. S., & Ravi, R. (2022). Pilot scale study on UV-C inactivation of bacterial endospores and virus particles in whole milk: evaluation of system efficiency and product quality. bioRxiv: The Preprint Server for Biology, 1-35.

Wilson, B. R., Roessler, P. F., Van Dellen, E., Abbaszadegan, M., & Gerba, C. P. (1992, Nov 15–19). Coliphage MS2 as a UV water disinfection efficacy test surrogate for bacterial and viral pathogens Proceedings of the American Water Works Association Water Quality Technology Conference, Toronto, Canada.

Wu, D., You, H., Jin, D., & Li, X. (2011). Enhanced inactivation of Escherichia coli with Ag- coated TiO2 thin film under UV-C irradiation. Journal of Photochemistry and Photobiology A: Chemistry, 217(1), 177–183.

Yaun, B. R., Sumner, S. S., Eifert, J. D., & Marcy, J. E. (2003). Response of Salmonella and Escherichia coli O157:H7 to UV energy. Journal of Food Protection, 66(6), 1071–1073.

